# Depletion of collagen IX alpha2 interferes osteochondral homeostasis of the knee joint which ultimately causing osteoarthritis-like articular cartilage damage

**DOI:** 10.1101/2022.09.24.509343

**Authors:** Rui Dong, Huihui Xu, Pinger Wang, Liang Fang, Luwei Xiao, Shuaijie Lv, Peijian Tong, Hongting Jin

## Abstract

As one of the branched chains of Type IX collagen (Col9), Collagen IX alpha2 (Col9α2) has been reported to be associated with several orthopedic conditions. To probe the relationship between Col9α2 and knee osteoarthritis (KOA), we performed a systematic analysis of Col9α2-deficient (Col9α^-/-^) mice using whole-mount skeletal staining, Micro-CT (μCT), biomechanics, histomorphometry, immunohistochemistry (IHC), immunofluorescence (IF) and Enzyme-linked immunosorbent (Elisa). Although whole-mount skeletal staining displayed no difference in bone length and ossification between Col9α^-/-^ mice and wild-type (Col9α2^+/+^) mice at mid-gestation and adult stages, the knee joint exhibited dramatic discrepancies. Specifically, the subchondral bone (SCB) in the knee joint of Col9α^-/-^ mice became sparse and deformed in the early stage, with altered bone morphometric parameters, reduced load-bearing capacity, dysfunctional bone homeostasis (decreased osteogenesis capacity and elevated bone resorption capacity), diminished cartilage proteoglycans and disrupted cartilage extracellular matrix (ECM) anabolism and catabolism compared with the Col9α2^+/+^ mice. In the late stage, the cartilage degeneration in Col9α2^-/-^ mice were particularly pronounced compared to Col9α2^+/+^ mice, as evidenced by severe cartilage destruction and a marked reduction in cartilage thickness and area. Overall, Col9α2 is essential for maintaining osteochondral homeostasis in the knee joint of mice, and the absence of this gene is accompanied by distinct sclerosis of the SCB and a reduction in load-bearing capacity; in the late stage, in the lack of SCB stress inhibition, excessive load is consistently exerted on the cartilage, ultimately leading to osteoarthritic-like articular cartilage damage. Hence, Col9α2 may serve as a potential candidate biomarker associated with KOA.

## Introduction

Type IX collagen (Col9) is heterotrimeric collagen consisting of three different α chains (α1, α2 and α3), expressed mainly in cartilage and found on the surface of tissues containing type II collagen (Col2), such as articular cartilage and vitreous humor [1, 2]. As fiber-related collagen with an interrupted triple helix, Col9 assembles with Col2 to form heterogeneous fibers. It cross-links with Col2 located on the fibril surface, with its non-collagenous NC4 structural domain projecting outwards [3]. Multiple interactions with other cartilage matrix proteins have been assigned to this structural domain, suggesting that Col9 bridges the collagen network and other extracellular matrix (ECM) superstructures [4].

As an integral component of articular cartilage and bone matrix [5–7], polymorphisms in Col9 have been reported to be associated with several orthopedic conditions. As one of the branched chains of Col9, Collagen IX alpha2 (Col9α2) has been most frequently reported to be involved in lumbar disc degeneration [8–10] and lumbar spinal stenosis [6, 11]. Our previous study has also demonstrated that the absence of Col9α2 leads to osteochondral remodeling of the cartilage endplate and inhibits ECM synthesis, which accelerates matrix degradation and chondrocyte hypertrophy in the intervertebral disc tissue [7]. However, some studies have also found a strong relationship between Col9α2 and the development of osteoarthritis (OA) of the hip [12] and multiple epiphyseal dysplasia [13, 14].

A recent study suggested that Col9α2 may be a potential candidate biomarker associated with OA, based on a comprehensive systematic analysis that reported a positive relationship between Col9α2 and OA [15], and combined with the strong correlation of Col9α2 with orthopedic diseases, it is reasonable to speculate that deletion of the Col9α2 is also implicated in the development of knee osteoarthritis (KOA). What are the specific changes and the underlying mechanisms if this is the case? Therefore, based on the above hypothesis, the current study provides the first evidence for the contribution of Col9α2 to osteochondral homeostasis in the knee joint and further observed the correlation with time.

## Materials and Methods

### Mice

Collagen IX alpha2-deficient (Col9α2^-/-^) mice were constructed from the Nanjing Biomedical Research Institute of Nanjing University (Grade SPF II, XM002445). Briefly, chimeric mice (Col9α2^+/-^) were generated and mated them for 10 generations. Col9α2^-/-^ mice and their wild-type (Col9α2^+/+^) littermates were applied for all experiments. All animals were housed in constant temperature (22°C) and humidity (40±5%), pathogen-free room, exposed to a controlled light/dark cycle (12h/12h) with solid rodent chow and water ad libitum. All studies were approved by the Committee on the Ethics of Animal Experiments of Zhejiang Chinese Medical University (Hangzhou, China; approval no. 20190401-10).

### Whole-mount skeletal staining

Skeletal staining of E18.5 and P28 mice were performed with 0.03% Alcian blue (Sigma-Aldrich, A5533) and 0.005% Alizarin red (Sigma-Aldrich, A3157) solutions [16]. These samples were hyalinized in a graded sequence of glycerol and potassium hydroxide and finally stored in 100% glycerol at 4°C for imaging on a C-DSD230 stereomicroscope (Nikon, Japan).

### Tissue sample and blood sample preparation

All mice were sacrificed at specific time points and the knee joint samples were surgically harvested; specimens were fixed in 4% paraformaldehyde for 48h, decalcified in 14% EDTA solution (PH=7.4) for approximately 14 days, then dehydrated and embedded in paraffin, stored at room temperature for further use.

The arterial blood of mice was collected by ophthalmectomy. EP tubes containing blood samples were left at room temperature for 30min, centrifuged at 3000 rpm/min for 15 min at 4°C, the serum was isolated and stored at −80°C pending use.

### Micro-CT Analysis (μCT)

The representative radiographic images of the knee joint samples were collected by micro-CT equipment (Skyscan 1176, Bruker, Kontich, Belgium). Briefly, the samples were scanned for approximately 400 slices at a resolution of 9μm. The region of interest (ROI) was defined between the proximal tibia growth plate and the tibial plateau, and the three-dimensional (3D) structure and ROI of the knee joint samples were reconstructed by using NRecon software (Bruker, Belgium). Several indexes were evaluated as follow: (1) bone mineral density (BMD, g/cm^3^); (2) bone volume fraction (BV/TV, %); (3) average trabecular thickness (Tb.Th, mm); (4) connectivity density (Conn.Dn); (5) average trabecular separation (Tb.Sp, mm).

### Mechanical test

To test the load-bearing capacity of the proximal tibia metaphysis of the samples, static loading was applied to the tibial plateau at 1 mm/min using an axial compression tester (EnduraTec TestBenchTM system, Bose Corp., Minnetonka, MN, USA). Subsequently, the appearance of the first mechanical turning point which attributed to specimen deformation, was defined as load-bearing capacity.

### Histochemistry and histomorphometry

Paraffin-embedded samples were cut into 3μm sagittal sections as described previously [17]. After deparaffinized in xylene and dehydrated in a series of graded ethanol, the sections were stained with Toluidine Blue and imaged on a light microscope (Axio Scope A1, ZEISS, Germany). The area and thickness of cartilage were calculated using OsteoMeasure Software (OsteoMetrics, Inc., Atlanta, GA, United States). Moreover, cartilage degeneration scores were determined by two blinded observers according to Osteoarthritis Research Society International (OARSI) scoring system as described previously [18].

### Tartrate-resistant acid phosphatase (TRAP) staining

After deparaffinized in xylene and dehydrated in a series of graded ethanol, the sections were stained with TRAP and counterstained with hematoxylin and imaged on a light microscope (Axio Scope A1, ZEISS, Germany), and the number of osteoclasts (N.Oc/T.A) was quantified as previously [19].

### Immunohistochemistry (IHC) and immunofluorescence (IF)

After endogenous peroxidase reduction, antigen retrieve and non-specific background elimination, sections were treated with anti-Col9α2 (diluted 1:100, Santa Cruz, sc398130, USA), anti-Col2 (diluted 1:1000, Millipore, mab1330, USA), anti-Runx2 (diluted 1:200, Abcam, ab76956, UK), anti-Mmp13(diluted 1:200, Abcam, ab39012, UK), anti-Col10 (diluted 1:1000, Abcam, ab58632, UK), anti-ALP (diluted 1:200, arigo, ARG57422, Taiwan, China), anti-Adamts5 (diluted 1:1000, Novus, NB100-74350, USA) primary antibody incubated overnight at 4 °C. For IHC staining, secondary goat anti-mouse/rabbit antibody (diluted 1:1000, Invitrogen, 31234, USA) was added for 20 min at room temperature on the second day. Positive staining of sections was presented by diaminobenzidine solution (ZSGB-bio, ZLI-9017, Beijing, China) while hematoxylin for counterstaining and imaged on a light microscope (Axio Scope A1, ZEISS, Germany). For IF staining, the sections were incubated with fluorophore-conjugated secondary antibody at room temperature for 20 min without the light, and then the sections were counterstained with DAPI and observed under a fluorescence microscope (Carl Zeiss AG). Finally, images were quantitatively analyzed by Image-Pro Plus 6.0 (Media Cybernetics, Rockville, Maryland, USA).

### Enzyme linked immunosorbent (Elisa) assay

Serum concentrations of bone turnover markers β-isomerized C-terminal telopeptides (β-CTX) and Procollagen Type I intact N-terminal Propeptide (PINP) were determined using an Enzyme-linked immunosorbent assay (Elisa) kits according to manufacturer’s procedures (Jiancheng bioengineering, H285 and H287, Nanjing, China). The absorbance readings were made at 450nm using a spectrophotometer (BioTek ELx800), and the concentrations of samples were calculated from the standard curve.

### Statistical analysis

All data were described as mean ± SD. One-way ANOVA test and independent-samples t-test, which was used in the present research, were performed using SPSS software (version 24.0, IBM Corporation, NY, USA). *P*<0.05 was considered to be statistically significant (*), meanwhile *P*<0.01 was classified as highly significant (#).

## Results

### Col9α2 depletion showed no difference in skeleton morphology in mice

Theoretically, the expression of Col9α2 protein may decrease sharply after Col9α2 gene depletion. As shown in Figure 1A, Col9α2 immunofluorescence appeared that the protein was mainly expressed in articular cartilage and the medullary cavity of subchondral bone (SCB), the expression content of SCB was relatively higher. As we expected, the expression of Col9α2 was significantly decreased in Col9α2^-/-^ mice. To figure out whether Col9α2 deficiency could interfere with the skeleton development in mice, we then conducted whole-mount skeletal staining by using embryos (E18.5) and adult mice (P28). The results displayed no difference in bone length and ossification between knockout and Col9α2^+/+^ mice at mid-gestation and adult stages (Figure 1B).

**Figure 1.**
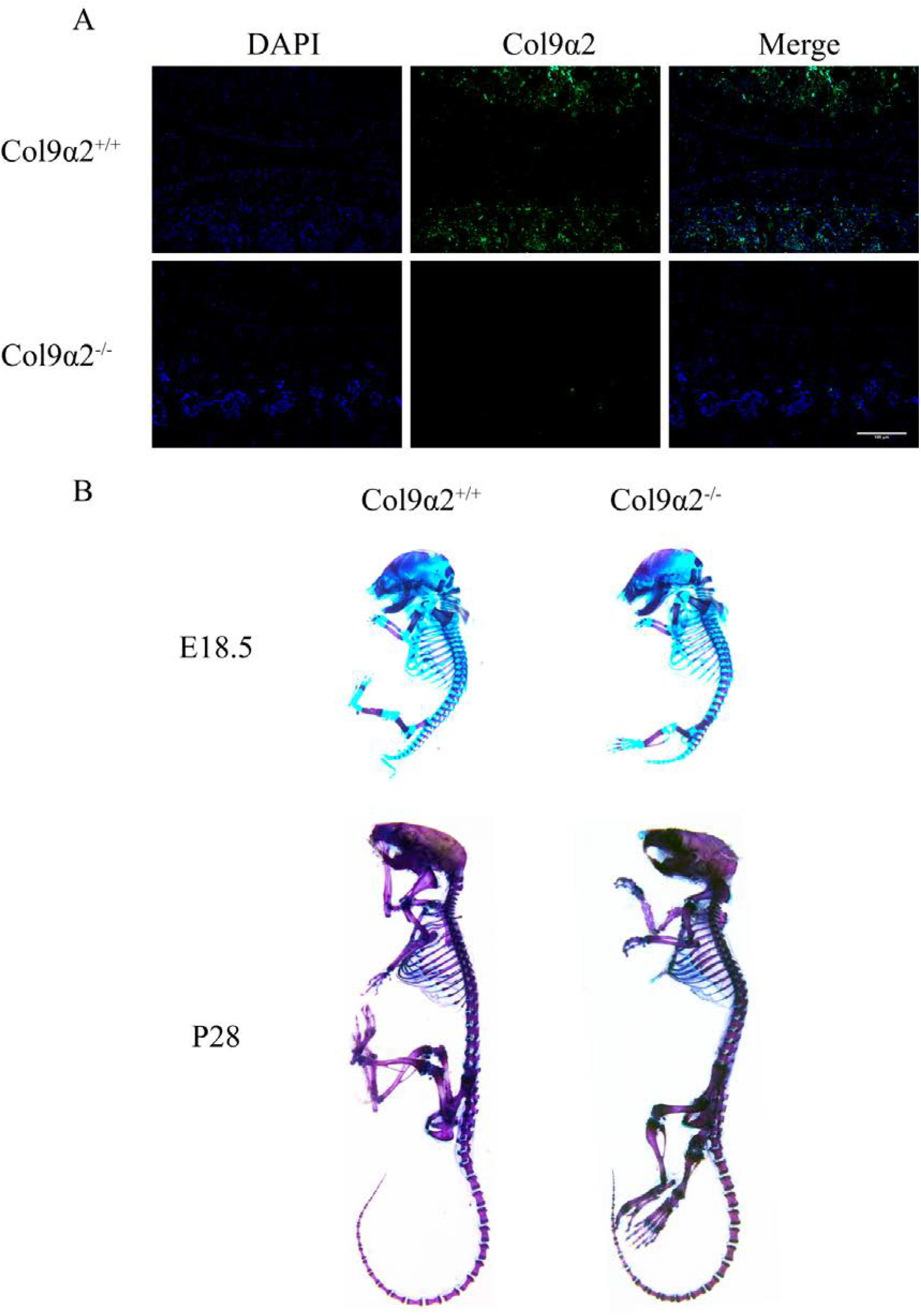
Immunofluorescence of Col9a2 and whole-mount skeletal staining of mice. A Immunofluorescence of col9a2 in 4-week-old mice. B Whole-mount skeleton staining of embryos (E18.5) and adult mice (P28).

### Col9α2 deficiency disrupts the homeostasis of subchondral bone by activating osteoclasts and inhibiting osteoblasts

Even though, as mentioned, there were no differences in skeletal appearance between the two types of mice, we wanted to find out if there were any clues to abnormalities in the microstructure of the skeleton. As shown in Figure 2A, 3D reconstruction analysis of μCT scanning exhibited that the microstructure of the tibial plateau SCB in Col9α2^-/-^ mice was thinner and sparser than that of Col9α2^+/+^ mice at 4w, 8w and 12w. Morphometric analysis of μCT data showed that BMD, BV/TV, Tb.Th and Conn.Dn were significantly lower in the corresponding regions of the Col9α2^-/-^ mice at 4w, 8w and 12w compared to Col9α2^+/+^ mice (Figure 2B-E), whereas Tb.Sp was considerably higher (Figure 2F).

**Figure 2.**
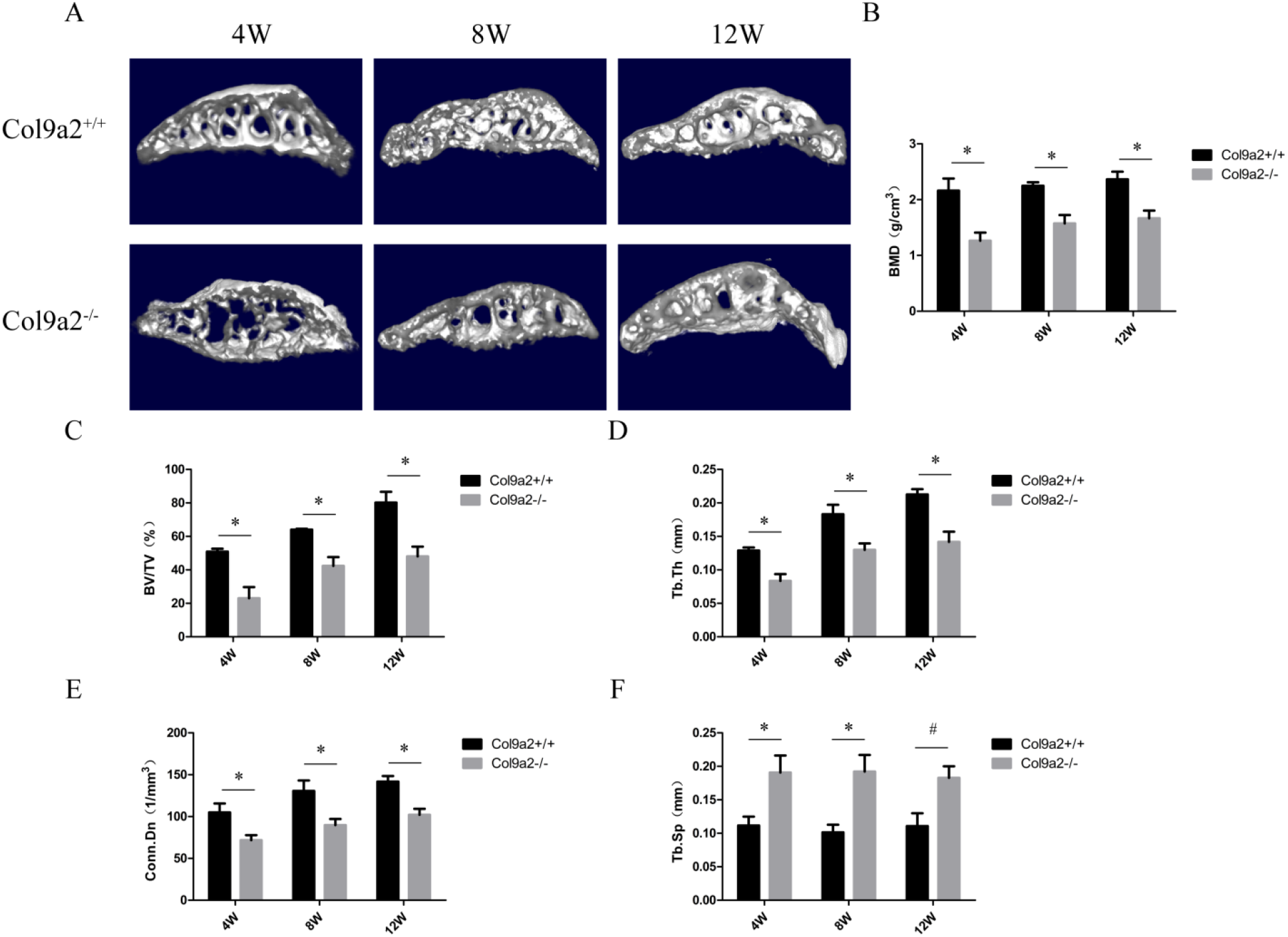
Image and morphometric analysis of μCT of early-stage Col9α2^-/-^ mice. A 3D reconstruction of the subchondral bone of the knee joint. B-E Morphometric analysis of μCT, including BMD (B); BV/TV (C); Tb.Th (D); Conn.Dn (E) and Tb.Sp (F). All data were expressed as mean ± SD, \**P* < 0.05, #*P* < 0.01.

To further explain these findings, we then discuss them by histopathology, biomechanics and serology. Toluidine blue staining manifested that the articular cartilage of Col9α2^-/-^ mice had fewer proteoglycans at 4w, 8w and 12w, although there was no apparent change in thickness. Notably, the SCB of Col9α2^-/-^ mice was considerably deformed compared to Col9α2^+/+^ mice (Figure 3A). To clarify whether these changes could affect the mechanical properties of the SCB, we examined the load-bearing capacity of the proximal tibial metaphysis. As expected, specimens from Col9α2^-/-^ mice were more prone to deformation under the same mechanical loading (Figure 3B). PINP and β-CTX are important reference indicators for evaluating osteogenesis and bone resorption, respectively, and in agreement with the imaging results, Col9α2^-/-^ mice also down-regulated serum PINP concentration and up-regulated serum β-CTX concentration (Figure 3C, D), reflecting the possibility of reduced osteogenesis and increased bone resorption in Col9α2^-/-^ mice. To verify this hypothesis, we then performed TRAP staining and IHC for osteogenesis-related proteins (ALP and Runx2), which demonstrated that Col9α2^-/-^ mice had a dramatically higher capacity for bone resorption compared to Col9α2^+/+^ mice, as well as a remarkably lower capacity for osteogenesis (Figure 4A-F).

**Figure 3.**
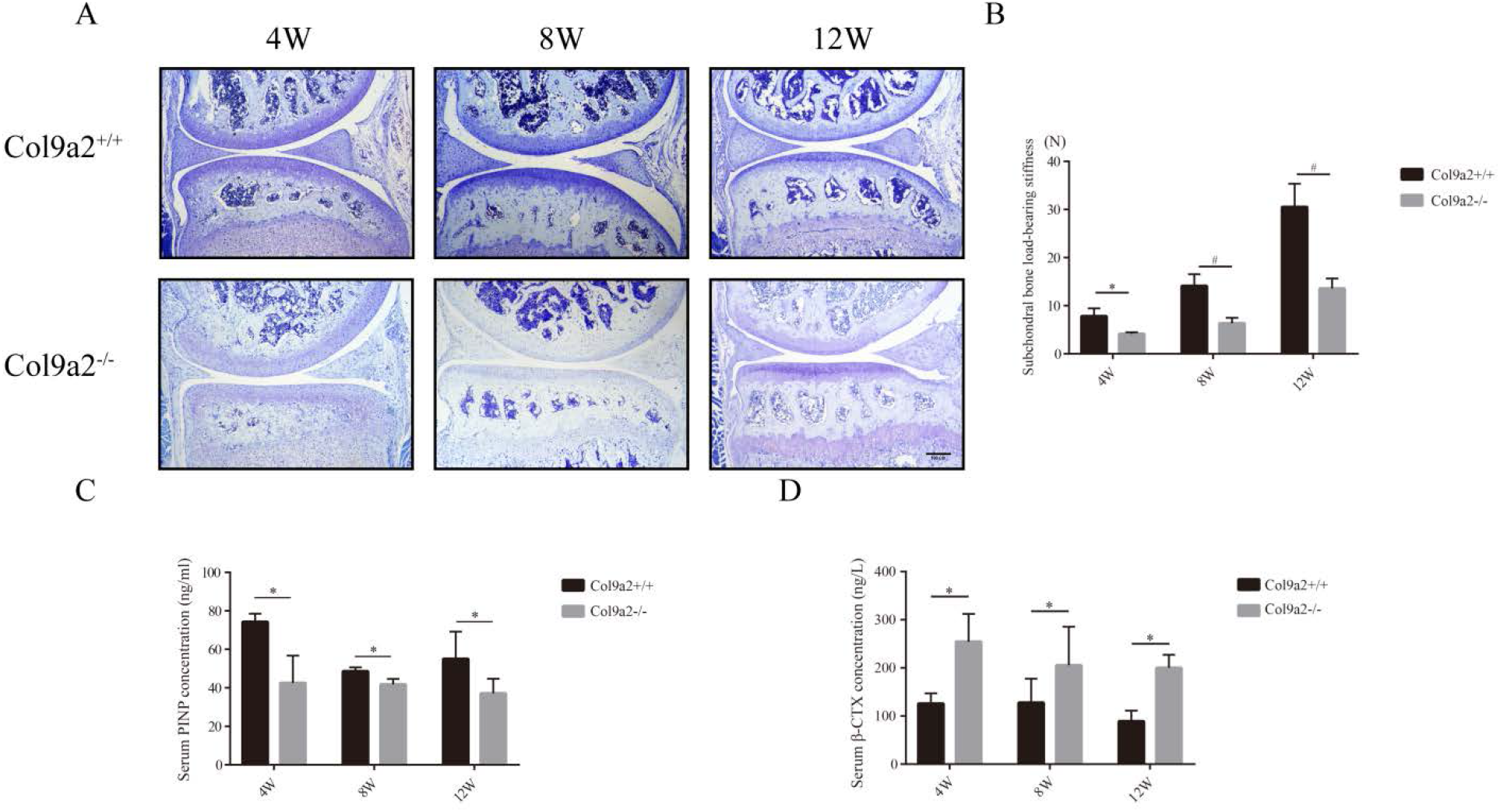
Histopathological, biomechanical and serological changes of early-stage Col9α2^-/-^ mice. A Toluidine blue staining. B Load-bearing capacity of the proximal tibia metaphysis. C,D Serum concentrations of β-CTX (C) and PINP (D). All data were expressed as mean ± SD, \**P* < 0.05, ^#^*P* < 0.01.

**Figure 4.**
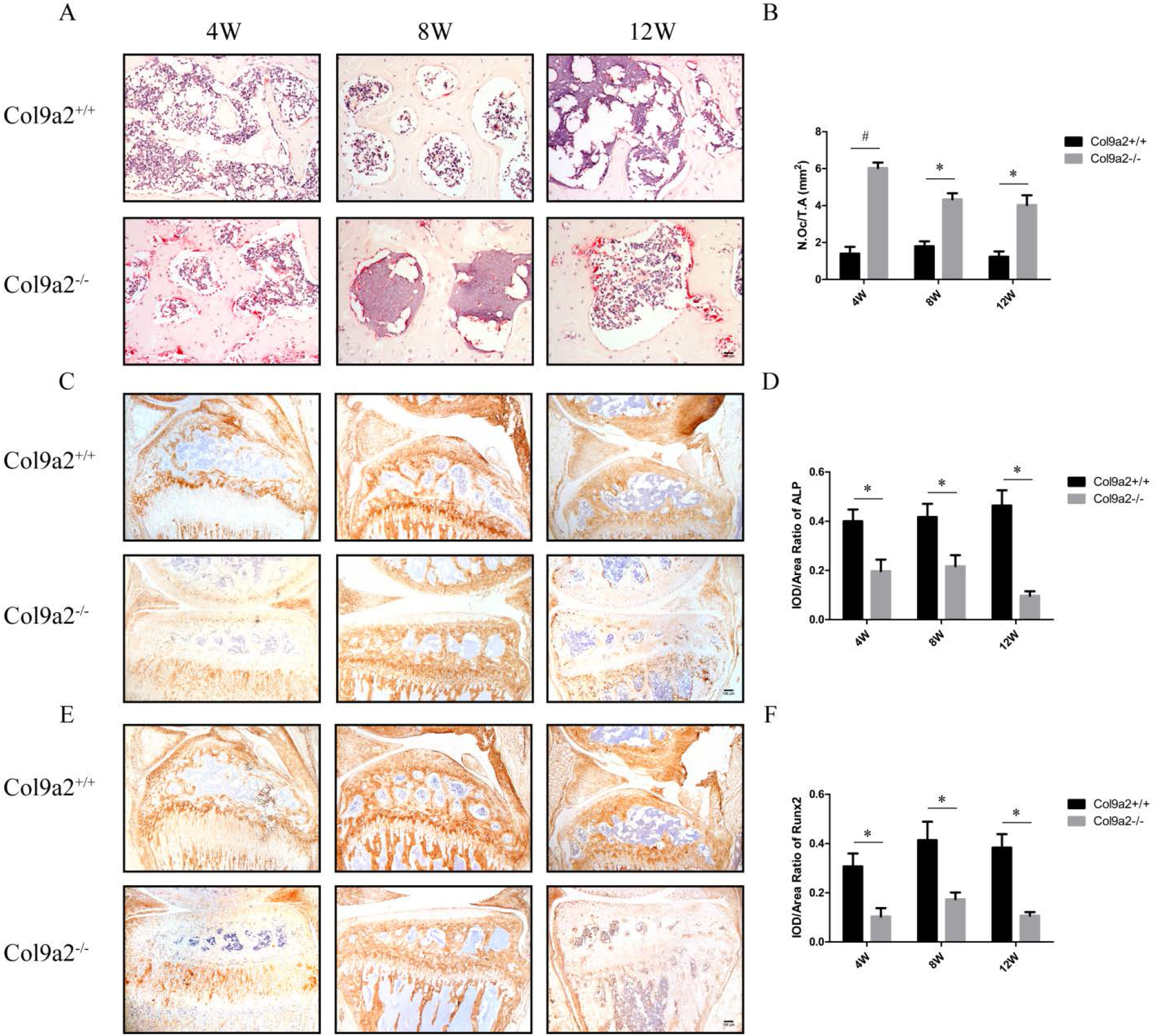
TRAP staining and IHC for osteogenic-related proteins of early-stage Col9α2^-/-^ mice. A,B TRAP staining (A) and osteoclastometric analysis (B). C,D IHC (C) and metrological analysis (D) of ALP. E,F IHC (E) and metrological analysis (F) of Runx2. All data were expressed as mean ± SD, \**P* < 0.05, #*P* < 0.01.

### Col9α2 depletion led to early articular cartilage degeneration and finally caused osteoarthritis-like articular cartilage damage

Articular cartilage and SCB seem like a functional unit, the underlying bone plate is highly vascularized and provide nutrition for upper hyaline cartilage. We believe that when SCB goes abnormal, cartilage can’t stay out of it. To testify our hypothesis, we conducted further studies.

As we described above, in addition to significant deformation of the SCB, there was a reduction in proteoglycans compared to Col9α2^+/+^ mice, although there was no marked change in articular cartilage thickness in Col9α2^-/-^ mice at 4w, 8w and 12w (Figure 3A). To explain the changes identified by histology, we performed IHC. IHC of the major components of the cartilage matrix (Col2 and Aggrecan) illustrated the expression of both proteins in the ECM of cartilage was dramatically reduced in the Col9α2^-/-^ mice compared to Col9α2^+/+^ mice (Figure 5A-D), in other words, the cartilage in Col9α2^-/-^ mice degraded faster. Since cartilage anabolism and catabolism are tightly linked, we then performed IHC on proteins related to cartilage catabolism. Among the chondrolysis-related proteins, Col10 is involved in the terminal differentiation of chondrocytes, Adamts5 is responsible for the degradation of Aggrecan, and Mmp13 is capable of degrading Col2 in the cartilage matrix. Surprisingly, whether Col10, Adamts5 or Mmp13, the expression of these proteins was substantially increased in Col9α2^-/-^ mice compared to Col9α2^+/+^ mice, demonstrating ECM degradation (Figure 6A-F). These results insisted that Col9α2 deficiency could interfere with cartilage ECM anabolism and catabolism.

**Figure 5.**
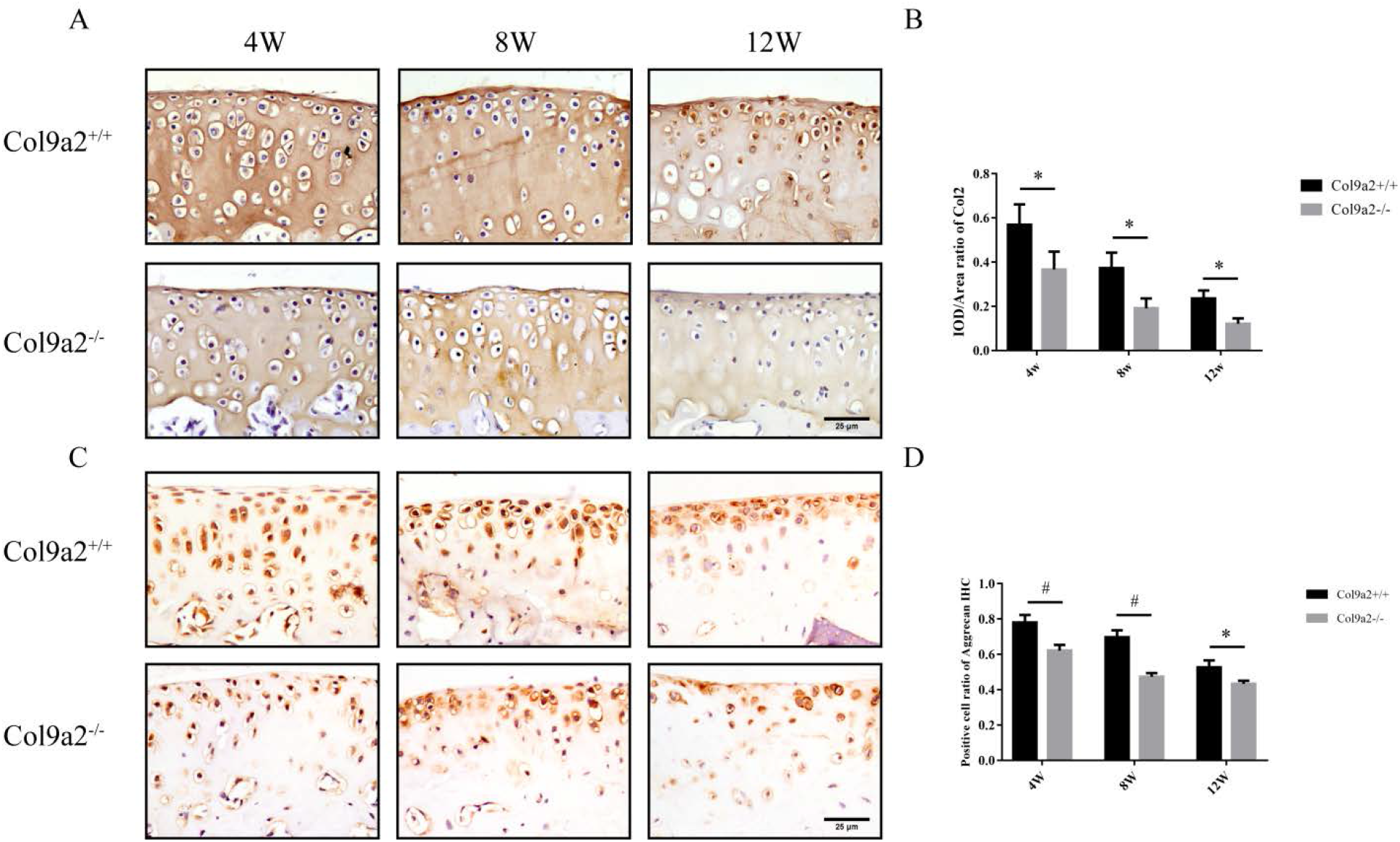
IHC of cartilage ECM anabolic-related proteins of early-stage Col9α2^-/-^ mice. A,B IHC (A) and metrological analysis (B) of Col2. C,D IHC (C) and metrological analysis (D) of Aggrecan. All data were expressed as mean ± SD, \**P* < 0.05, #*P* < 0.01.

**Figure 6.**
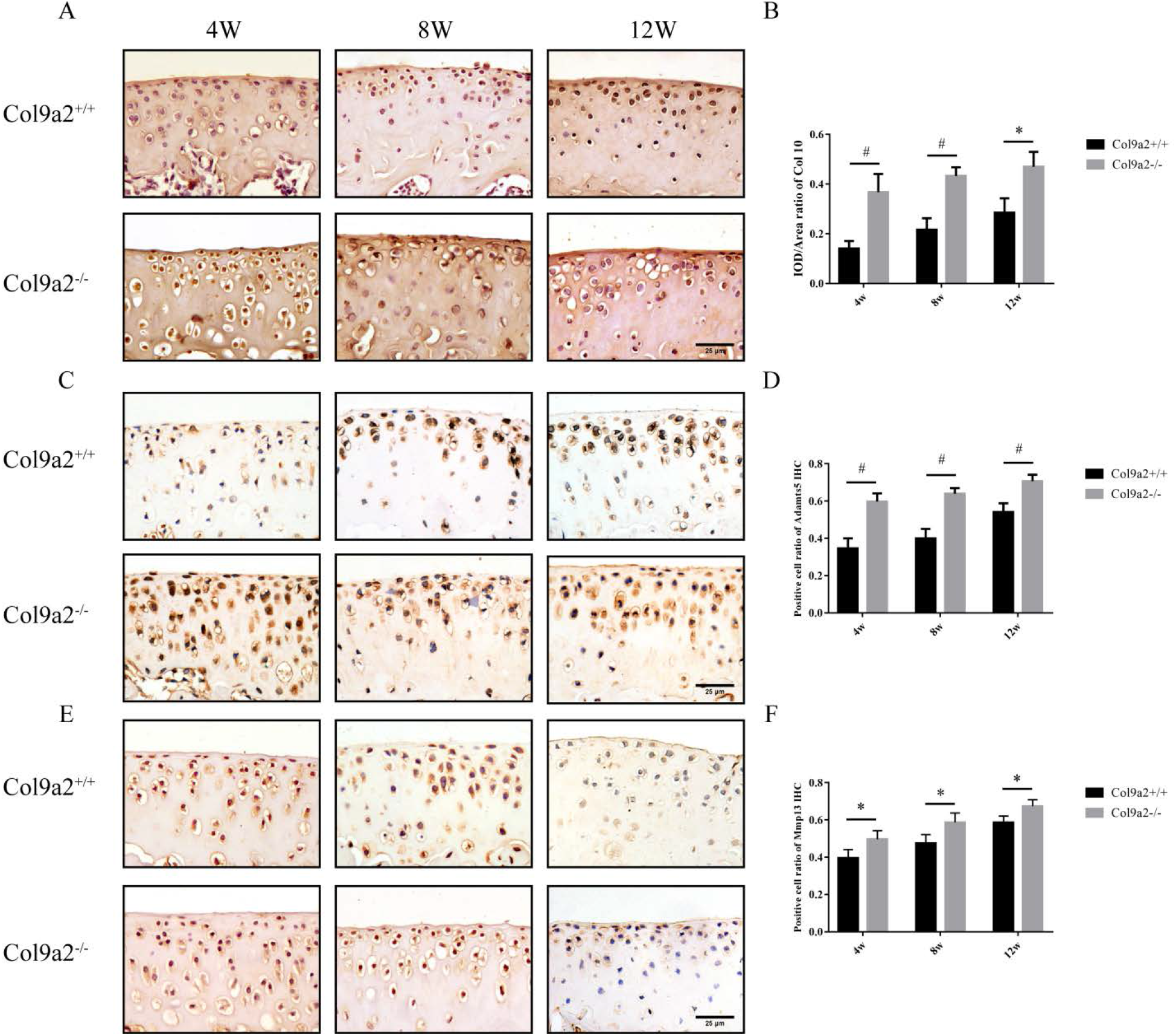
IHC of cartilage ECM catabolic-related proteins of early-stage Col9α2^-/-^ mice. A,B IHC (A) and metrological analysis (B) of Col10. C,D IHC (C) and metrological analysis (D) of Adamts5. E,F IHC (E) and metrological analysis (F) of Mmp13. All data were expressed as mean ± SD, \**P* < 0.05, #*P* < 0.01.

Based on the imbalance in osteochondral homeostasis in Col9α2^-/-^ mice, we hypothesized that Col9α2 deficiency would eventually cause the destruction of articular cartilage. To examine this hypothesis, we extended the observation time to 36w. Toluidine blue staining revealed that at 24w of age, the Col9α^-/-^ mice exhibited a marked narrowing of the joint space, reduced cartilage matrix staining and pronounced SCB sclerosis compared to Col9α2^+/+^ mice. At 36w of age, the knee joint degeneration in Col9α2^-/-^ mice was further aggravated with severe cartilage destruction, particularly on the tibial side, and the sclerosis of SCB was further exacerbated (Figure 7A). Bone morphometrics demonstrated that both the cartilage thickness and area of the tibial plateau were dramatically lower than in the Col9α2^+/+^ mice (Figure 7B, C). The OARSI of the knee joint was markedly higher in Col9α2^-/-^ mice than in the Col9α2^+/+^ mice (Figure 7D). The results of the 3D reconstruction analysis of μCT scans further confirmed the degeneration of the knee joint. 24 and 36 week old Col9α2^-/-^ knockout mice displayed distinct osteophytes growth with joint space narrowing compared to Col9α2^+/+^ mice (Figure 7E). However, morphometric analysis of μCT data showed no significant differences in BV/TV, Tb.Th and Tb.Sp parameters (Figure 7F-H).

**Figure 7.**
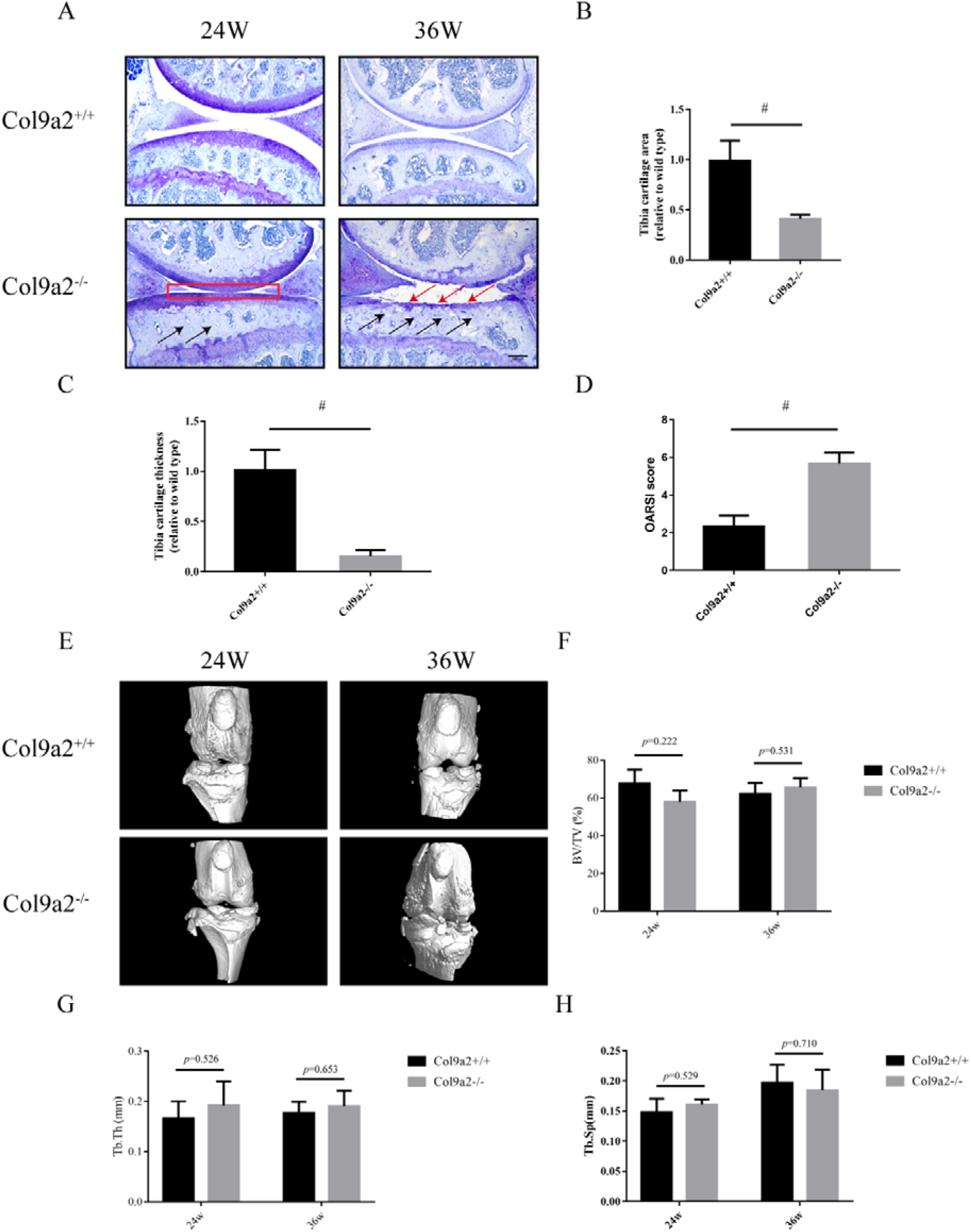
Histopathological staining and μCT analysis of late-stage Col9α2^-/-^ mice. A Toluidine blue staining. Red box indicates the narrowed joint space, black arrows indicate the area of subchondral osteosclerosis and red arrows indicate the area of cartilage defects. B,C Cartilage thickness (B) and area (C) of the tibial plateau. D OARSI score. E 3D reconstruction of the knee joint. F-H Morphometric analysis of μCT, including BV/TV (F); Tb.Th (G) and Tb.Sp (H). All data were expressed as mean ± SD, ^#^*P* < 0.01.

## Discussion

The progressive degeneration of joints inevitably occurs over the course of a lifetime as a consequence of natural aging or external damage. These are the major risk factors for the progression to osteoarthritis (OA), the most common degenerative bone and joint disease affecting approximately 250 million people worldwide [20–22]. The joint consists primarily of cartilage and SCB, separated by an osteochondral interface, which comprises the deeply packed cartilage and the underlying SCB. These individual components collaborate with each other to form a complex functional unit [23]. Anatomically, the articular cartilage is attached to the SCB, which is a richly vascularized tissue that provides nutrients and material exchange to the hyaline cartilage above [24]. Biomechanically, the stability and integrity of the articular cartilage are dependent on its interaction with the SCB [25], which provides mechanical support to the overlying articular cartilage during weight-bearing joint movements and is in a constant process of deformation and remodeling in order to adapt to the changing mechanical environment [26]. Thus, osteochondral homeostasis is essential for the maintenance of knee joint movement and function.

The collagen plays a pivotal role in osteochondral homeostasis, which provides the structural support for bone and cartilage, and their normal distribution in time and space ensures the normal morphology of bone and cartilage and their stable biomechanical properties such as resistance to compression and rotation [27]. Col9, together with Col2, forms the scaffold for the ECM in cartilage and is an important component of the collagen network within cartilage [28]. As one of the branched chains of Col9, Col9α2 has been shown to be associated with orthopedic conditions such as intervertebral disc degeneration and multiple epiphyseal dysplasia [6, 8–11], but the relationship with OA remains to be elucidated. In the present study, we first demonstrated by whole skeleton staining that the absence of Col9α2 could not affect skeletal development in mice. We then focused on the knee joint to observe the effect of Col9α2 deletion on osteochondral homeostasis. As expected, μCT results revealed remarkable changes in the microstructure and trabecular parameters of the tibial plateau SCB in early Col9α2^-/-^ mice compared to wild-type mice, which were confirmed by pathological staining and biomechanical tests. Bone homeostasis is a state in which bone resorption and bone formation are in a dynamic balance [29, 30]. Based on the marked alterations in SCB, we speculated that this might be due to an imbalance in bone homeostasis. Fortunately, Elisa, TRAP staining and IHC results displayed that Col9α2^-/-^ mice exhibited active osteoclastic function, whereas osteogenic function was reduced.

However, the degeneration of the cartilage apparently lagged behind the SCB, with no change in cartilage thickness, except for a decrease in proteoglycan, suggesting that the SCB in Col9α2^-/-^ mice were drastically altered than the cartilage in the early stages. Based on the strong association of SCB with cartilage, it is reasonable to assume that cartilage is altered in addition to a reduction in proteoglycan. IHC results demonstrated that Col9α2 deficiency interferes with the anabolic and catabolic metabolism of cartilage ECM, leading to later cartilage degeneration. Given this striking phenomenon, we are interested in further observing the effect of the absence of Col9α2 on osteochondral homeostasis of the knee joint at a later stage. μCT and pathological staining revealed that in addition to further subchondral osteosclerosis, the Col9α2^-/-^ mice had severely damaged cartilage with highly uneven surfaces, marked fissures and remarkably increased osteophytes growth and joint space narrowing compared to wild-type mice, as well as notably higher OARSI scores. However, surprisingly, the morphometric analysis of μCT indicated that there was no significant difference between the two groups of mice in terms of BMD, BV/TV, Tb.Th and Tb.Sp. We speculate that this result may be due to the early onset of osteogenesis and osteoclastic activity of Col9α2 deletion, resulting in sparse bone trabeculae and reduced BMD in the SCB of the knee, although osteoarthritic-like changes in the knee were observed later, it was not at the same level as in wild-type mice and therefore was not comparable in the later stages. Therefore, the lack of statistical significance of the relevant parameters is not sufficient to negate the fact of sclerosis.

## Conclusion

In conclusion, Col9α2 is essential for the maintenance of osteochondral homeostasis in the knee joint of mice. After the knockout of this gene, the sclerosis of SCB is dramatically accompanied by the reduction of load-bearing capacity; in the late stages, excessive load is continuously applied to the cartilage in the presence of SCB stress depression, ultimately causing osteoarthritis-like articular cartilage damage.

## Acknowledges

We appreciate the great help/technical support/experimental support from the Public Platform of Medical Research Center, Academy of Chinese Medical Science, Zhejiang Chinese Medical University.

## Contributions

L Xiao, S Lv, P Tong and H Jin conceived of the studies and planned the experimental design. R Dong, H Xu, P Wang, L Fang performed the experiments and analyzed the data. H Xu wrote the manuscript, R Dong assisted with data interpretation and performed critical revisions on the manuscript. All authors approved the final manuscript.

## References

1. Paassilta P, Pihlajamaa T, Annunen S, Brewton RG, Wood BM, Johnson CC, et al. Complete sequence of the 23-kilobase human COL9A3 gene. Detection of Gly-X-Y triplet deletions that represent neutral variants. J Biol Chem. 1999; 274: 22469–75.

2. Pihlajamaa T, Vuoristo MM, Annunen S, Perala M, Prockop DJ, Ala-Kokko L. Human COL9A1 and COL9A2 genes. Two genes of 90 and 15 kb code for similar polypeptides of the same collagen molecule. Matrix Biol. 1998; 17: 237–41.

3. Eyre DR, Pietka T, Weis MA, Wu JJ. Covalent cross-linking of the NC1 domain of collagen type IX to collagen type II in cartilage. J Biol Chem. 2004; 279: 2568–74.

4. Bruckner P. Suprastructures of extracellular matrices: paradigms of functions controlled by aggregates rather than molecules. Cell Tissue Res. 2010; 339: 7–18.

5. Roberts S, Menage J, Duance V, Wotton S, Ayad S. 1991 Volvo Award in basic sciences. Collagen types around the cells of the intervertebral disc and cartilage end plate: an immunolocalization study. Spine (Phila Pa 1976). 1991; 16: 1030–8.

6. Hyun SJ, Park BG, Rhim SC, Bae CW, Lee JK, Roh SW, et al. A haplotype at the COL9A2 gene locus contributes to the genetic risk for lumbar spinal stenosis in the Korean population. Spine (Phila Pa 1976). 2011; 36: 1273–8.

7. Xu H, Dong R, Zeng Q, Fang L, Ge Q, Xia C, et al. Col9a2 gene deletion accelerates the degeneration of intervertebral discs. Exp Ther Med. 2022; 23: 207.

8. Annunen S, Paassilta P, Lohiniva J, Perala M, Pihlajamaa T, Karppinen J, et al. An allele of COL9A2 associated with intervertebral disc disease. Science. 1999; 285: 409–12.

9. Jim JJ, Noponen-Hietala N, Cheung KM, Ott J, Karppinen J, Sahraravand A, et al. The TRP2 allele of COL9A2 is an age-dependent risk factor for the development and severity of intervertebral disc degeneration. Spine (Phila Pa 1976). 2005; 30: 2735–42.

10. Aladin DM, Cheung KM, Chan D, Yee AF, Jim JJ, Luk KD, et al. Expression of the Trp2 allele of COL9A2 is associated with alterations in the mechanical properties of human intervertebral discs. Spine (Phila Pa 1976). 2007; 32: 2820–6.

11. Noponen-Hietala N, Kyllonen E, Mannikko M, Ilkko E, Karppinen J, Ott J, et al. Sequence variations in the collagen IX and XI genes are associated with degenerative lumbar spinal stenosis. Ann Rheum Dis. 2003; 62: 1208–14.

12. Nakki A, Videman T, Kujala UM, Suhonen M, Mannikko M, Peltonen L, et al. Candidate gene association study of magnetic resonance imaging-based hip osteoarthritis (OA): evidence for COL9A2 gene as a common predisposing factor for hip OA and lumbar disc degeneration. J Rheumatol. 2011; 38: 747–52.

13. Spayde EC, Joshi AP, Wilcox WR, Briggs M, Cohn DH, Olsen BR. Exon skipping mutation in the COL9A2 gene in a family with multiple epiphyseal dysplasia. Matrix Biol. 2000; 19: 121–8.

14. Muragaki Y, Mariman EC, van Beersum SE, Perala M, van Mourik JB, Warman ML, et al. A mutation in COL9A2 causes multiple epiphyseal dysplasia (EDM2). Ann N Y Acad Sci. 1996; 785: 303–6.

15. Zhang R, Guo H, Yang X, Li Z, Zhang D, Li B, et al. Potential candidate biomarkers associated with osteoarthritis: Evidence from a comprehensive network and pathway analysis. J Cell Physiol. 2019; 234: 17433–43.

16. Rigueur D, Lyons KM. Whole-mount skeletal staining. Methods Mol Biol. 2014; 1130: 113–21.

17. Xu HH, Li SM, Fang L, Xia CJ, Zhang P, Xu R, et al. Platelet-rich plasma promotes bone formation, restrains adipogenesis and accelerates vascularization to relieve steroids-induced osteonecrosis of the femoral head. Platelets. 2020: 1–10.

18. Xu HH, Li SM, Xu R, Fang L, Xu H, Tong PJ. Predication of the underlying mechanism of Bushenhuoxue formula acting on knee osteoarthritis via network pharmacology-based analyses combined with experimental validation. J Ethnopharmacol. 2020; 263: 113217.

19. Zhang P, Xu H, Wang P, Dong R, Xia C, Shi Z, et al. Yougui pills exert osteoprotective effects on rabbit steroid-related osteonecrosis of the femoral head by activating beta-catenin. Biomed Pharmacother. 2019; 120: 109520.

20. Felson DT, Niu J, Yang T, Torner J, Lewis CE, Aliabadi P, et al. Physical activity, alignment and knee osteoarthritis: data from MOST and the OAI. Osteoarthritis Cartilage. 2013; 21: 789–95.

21. Sanchez-Adams J, Leddy HA, McNulty AL, O’Conor CJ, Guilak F. The mechanobiology of articular cartilage: bearing the burden of osteoarthritis. Curr Rheumatol Rep. 2014; 16: 451.

22. Hunter DJ, Bierma-Zeinstra S. Osteoarthritis. Lancet. 2019; 393: 1745–59.

23. Yuan XL, Meng HY, Wang YC, Peng J, Guo QY, Wang AY, et al. Bone-cartilage interface crosstalk in osteoarthritis: potential pathways and future therapeutic strategies. Osteoarthritis Cartilage. 2014; 22: 1077–89.

24. Imhof H, Sulzbacher I, Grampp S, Czerny C, Youssefzadeh S, Kainberger F. Subchondral bone and cartilage disease: a rediscovered functional unit. Invest Radiol. 2000; 35: 581–8.

25. Lories RJ, Luyten FP. The bone-cartilage unit in osteoarthritis. Nat Rev Rheumatol. 2011; 7: 43–9.

26. Madry H, van Dijk CN, Mueller-Gerbl M. The basic science of the subchondral bone. Knee Surg Sports Traumatol Arthrosc. 2010; 18: 419–33.

27. Ott SM. Bone strength: more than just bone density. Kidney Int. 2016; 89: 16–9.

28. Wu JJ, Woods PE, Eyre DR. Identification of cross-linking sites in bovine cartilage type IX collagen reveals an antiparallel type II-type IX molecular relationship and type IX to type IX bonding. J Biol Chem. 1992; 267: 23007–14.

29. Nakahama K. Cellular communications in bone homeostasis and repair. Cell Mol Life Sci. 2010; 67: 4001–9.

30. Eriksen EF. Cellular mechanisms of bone remodeling. Rev Endocr Metab Disord. 2010; 11: 219–27.

